# Mechanical constraints organize 3D tissues and orchestrate muscle differentiation

**DOI:** 10.1101/2024.10.03.616457

**Authors:** Irène Nagle, Lorijn van der Spek, Paul Gesenhues, Thierry Savy, Laurent Réa, Alain Richert, Mathieu Receveur, Florence Delort, Sabrina Batonnet-Pichon, Claire Wilhelm, Nathalie Luciani, Myriam Reffay

## Abstract

Biological tissues achieve proper shape and ordered structures during development through responses to internal and external signals, with mechanical cues playing a crucial role. These forces guide cellular organization, leading to complex self-organizing structures that are foundational to embryonic patterns. Emerging theories and experiments suggest that “topological morphogens” drive these processes. Despite the predominance of three-dimensional (3D) structures in biology, studying 3D tissues remains challenging due to limited model systems and the complexity of modeling. Here, we address these challenges by using self-organized cellular aggregates, specifically spindle-shaped C2C12 myoblasts, subjected to controlled mechanical stretching. Our findings reveal that these cells form a multilayered, actin-oriented tissue structure, where mechanical forces drive long-range 3D organization and muscle differentiation. Notably, tissue surface emerges as a hotspot for differentiation, correlating with directional order as shown by single molecule fluorescent *in situ* hybridization.

**Significance Statement:** We explore how cells work together to form complex structures, particularly in 3D, using muscle precursors cells (C2C12 myoblasts) as a model. By applying controlled stretching forces, we found that these cells self-organize into layered tissues that guide their transformation into muscle. This research highlights the critical role of physical forces in shaping tissues, suggesting that the way cells are physically arranged and stretched in three dimensions can significantly influence their behavior and function. Our findings offer new insights into how tissues develop and could have implications for tissue engineering, where creating the right 3D environment is key to successful tissue growth and repair.

---

**M**orphogenesis is a remarkable process because of its robustness across scales and conditions based on the integration of internal and external signals. Among these feedbacks, mechanical stimuli are of enormous importance for pattern emergence in multicellular systems and organisms (1–4). These signals are transduced by cellular mechanosensors and give rise to macroscopic rearrangements, supracellular movements and complex self-organizing structures that are at the birth of early embryonic patterns (5, 6). Developmental processes depend on a wide range of forces that correlate with morphogen gradients (7), but also with cell shape and local organization. How cells can work together as dynamic collective ensembles to achieve a proper organization and how an organism is shaped by mechanical forces during development are still crucial open questions.

The idea of “topological morphogens” (8) is beginning to emerge, meaning that biological functions could arise from topological mechanisms that create reproducible and robust conditions to control them (9–12). Indeed, topological similarities across scales can be found in nature. For example, the network of epithelial tubes providing transport within the body is reminiscent of intestinal arrangements and is related to folding and buckling processes (13). Thus, development is actually based on a relatively small number of key processes that the variability of tissues and organisms may have hidden at first glance. Theories of out-of-equilibrium active matter provide a new general framework to study these phenomena in multicellular systems (14, 15). Dimensionality (16), curvature (17, 18) and topological defects (19–21) seem to be the main ingredients of the interplay between topology and biological function.

The emergence of a third dimension in cell monolayers is, for example, specifically related to defect dynamics at different scales and in different systems, whether for extrusions (22) or for multicellular protrusions in confined myoblasts (23). Conversely, we might wonder whether the emergence of a three-dimensional shape can help to organize cells and subsequently improve their differentiation. More generally, the role of orientational order and anisotropy in biological processes (24) and in model cell monolayers (25) is beginning to be taken into account in a more systematic way. In this context, myogenesis is of particular interest because muscle function is based on a multiscale architecture, from the actin filaments to the syncytial cells that make up the muscle and support its contractility.

In addition, while biological systems are mostly three-dimensional, studies of 3D tissues are still sparse, hampered by several obstacles. First, there is a lack of purely cellular model systems that can be constrained in a controlled manner. 3D mechanical constraints are generally applied using 3D scaffolds. The presence of the scaffold adds complexity, as the topology of the biomaterials and their adhesion to the substrate need to be considered (16). Secondly, increasing dimensionality opens up a wider range of possible configurations that may prove difficult to model. However, in nematic synthetic fluids, it gives rise to fascinating disclination loops reminiscent of topological defects in 2D (26). In addition, deciphering the interplay between 3D tissue formation and differentiation may be a key challenge for tissue engineering applications.

Here, we address this question and challenge by developing an approach based on self-organized purely cellular aggregates whose geometry is controlled using mechanical constraints. More specifically, we study how spindle-shaped C2C12 myoblast cells organize in 3D and differentiate in muscular cells when stretched. To this end, linear, uniaxial and high amplitude stretching is applied over a large tissue using a magnetic device first designed to drive embryonic stem cell differentiation towards the cardiac pathway (27). We show that C2C12 cells organize in an active tissue with a fascinating multilayered structure that drives actin orientation and muscle cell differentiation. Although the third dimension is of great importance in the orientation and the differentiation, applied mechanical forces actually drive the long-range ordering of the 3D tissue and its subsequent differentiation. Furthermore, the surface of this model tissue emerges as a hotspot in differentiation. Using single molecule fluorescent *in situ* hybridization, we demonstrate that the directional order is correlated with the obtained differentiation pattern. The application of quantitative approaches proves to be powerful to characterize patterns and mechanosensitivity in these model tissues.

## Results

### Stretching myoblasts in a magnetic stretcher device

To generate cohesive and mechanically stimulable 3D model tissues presenting possible orientational order, we form multicellular aggregates of C2C12 myoblasts (of approximatively 10^5^ cells) (Fig. 1a). C2C12 cells are muscle cell precursors (28), i.e. they can differentiate into myotubes after collective alignment, long-range orientation and fusion steps (29–31). The cells are first labelled with superparamagnetic nanoparticles (NP) to give them magnetic properties (32). Incorporation of citrated iron oxide NP does not change the metabolic activity or differentiation potential of C2C12 cells and is therefore biocompatible (Fig. S1 and S2b). Magnetically labelled cells are equally distributed on two face-to-face magnetic microattractors forming two domes of non-cohesive cells that are brought into contact on day 0. One of the two attractors is mobile to stimulate and stretch the formed 3D aggregate (Fig. 1a). Maximal near the soft iron tips (in the range of a few hundreds pN) and close to zero in the central part of the setup (Fig. S3a-b), the magnetic forces are smaller than the adhesion between cells once established or the pulling forces that cells can exert (33). They remain constant over the course of the experiment as the magnetization M is constant over a week (Fig. S3c). After nine hours in complete medium, a cohesive aggregate with a smooth surface is formed (Fig. 1b-c).

The medium is then changed to differentiation medium (2% horse serum) on day 1 (D1). Linear uniaxial stretching (1 µm /3 min for 6 hours) is started. The magnetic forces are not altered by moving the mobile micromagnet apart during stretching as can be seen from the force profiles (Fig. S3a-b).The magnets thus act as clamps to hold the cohesive aggregate once formed in combination with cell adhesion to glass slides. The magnetic stretcher impose mechanical stress to the 3D multicellular aggregate by controlling the movement of the micromagnet (27), it stimulates mechanically and stretch the 3D structure (Fig. 1d-f) to drive cell alignment and differentiation. The selected speed is slow enough to allow cells to rearrange between each step (34, 35). The magnetic stretching device was adapted to muscle cell differentiation with large tissues (approximately 400µm height and 500µm diameter) and to apply high mechanical constraint with a maximum stretch of 30 %. The stretching rate was optimised to improve cell alignment while preventing potential tissue rupture. The multicellular aggregate is then kept in the magnetic stretcher for 24 to 48 hours (day 2 (D2) and day 3 (D3)), depending on the measurement performed.

**Fig. 1.**
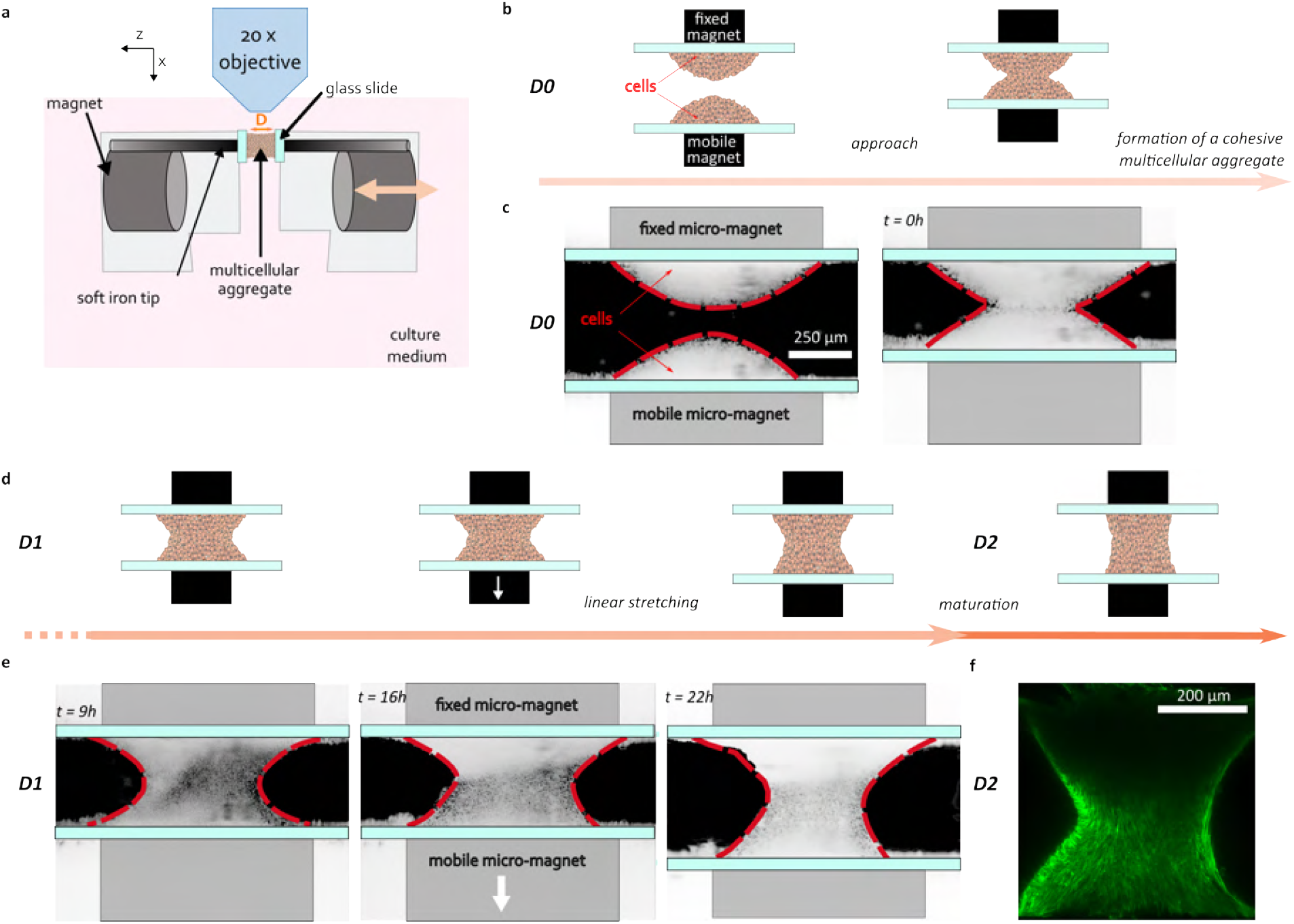
Stretching 3D model tissue. **a)** Schematic of the magnetic stretching system. A multicellular assembly of magnetic cells is supported by two micromagnets separated by a distance D. As one of the micromagnets is movable, the device imposes the mechanical stretching to the aggregate. **b-c)** Schematic and representative images of the formation of a cohesive multicellular aggregate in the magnetic stretcher. First, 5 · 10^4^ magnetic cells in suspension are placed on each micromagnet. The two pieces are then brought together and a cohesive aggregate is formed overnight. **d-e)** Schematic and representative images of the stretching of the multicellular aggregate and after one day of maturation in the magnetic stretcher. **f)** Representative 2-photon image of the stretched multicellular aggregate at a depth of 50 µm. Cells express LifeAct-GFP.

### Stretching aligns the cells

Actin, one of the major cellular proteins of the cytoskeleton, supports the overall structure of cells, and C2C12 cells are no exception. As early as day 1, multicellular aggregates show a collective organization. Interestingly, the orientation of the filaments at the surface of the aggregates in the magnetic stretcher is tilted almost perpendicular to the orientation in the core of the aggregate at a depth of 50 µm and below (Fig. 2 a-b).

**Fig. 2.**
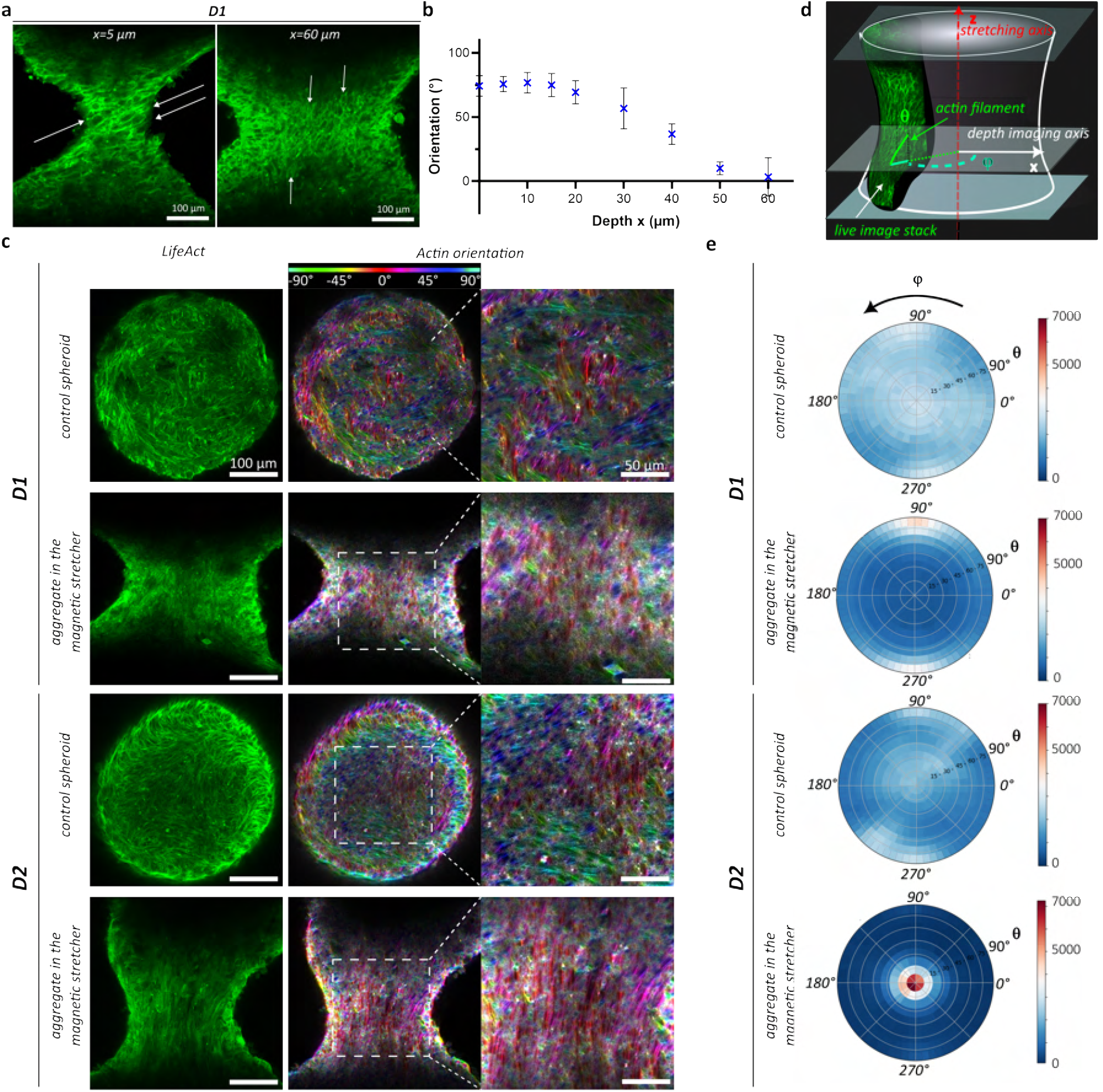
3D alignement of actin filaments driven by stretching. **a)** Representative 2-photon images of actin filaments (LifeAct-GFP) in C2C12 multicellular aggregate in the magnetic stretcher at 5 µm depth (left) and at 60 µm depth (right). Images were taken on D1 before stretching. At the surface, actin filaments (indicated by white arrows) are oriented with a tilt relative to the *z* stretching axis higher than 45^*°*^, whereas in the core of the aggregate they are parallel to the stretching axis. **b)** Orientation of actin filaments as a function of depth *x* for the aggregate in the magnetic stretcher on D1 before stretching (Mean +/-SD. N = 3). **c)** Left panels: representative images of C2C12 Lifeact-GFP control spheroids and stretched aggregates on D1 and D2 at approximately 50 µm depth. Actin is shown in green. Middle panels: corresponding images with filaments color-coded according to their projected orientation with respect to the stretching axis *z*. Right panels: zoom in on the central region of the color-coded orientation map. **d)** Schematic of the stretched aggregate to define the azimuthal angle *θ* with respect to the stretching axis *z* and the polar angle *ϕ* with respect to the depth imaging axis **e)** Stereoplot of the angular distribution of actin filaments measured between 0 and 120 *µ*m depth for control spheroids and for the stretched aggregates on D1 and D2. A peak in the direction of stretching is observed for the stretched aggregate at D2. N = 5 independent experiments for control spheroids and N = 4 independent experiments for aggregates in the magnetic stretcher.

This multilayered structure evolves when a mechanical stimulus is applied. At D1 and D2, actin filament orientation was imaged in aggregates formed in the magnetic stretcher and compared with control (CTL) 3D spheroids that were formed simultaneously with the stretched aggregates by magnetic moulding (36, 37) and left free of mechanical constraints in parallel under the same conditions (Fig. 2c). The curvature of these spheroids is chosen to be in the same range as the mean curvature of the stretched aggregates (210 ± 20 µm at D1).

In both systems, the number and the length of the actin filaments increase between day 1 and day 2 (Fig. 2c-e and Fig. S4), but are much greater in the stretched condition. In addition, the orientation of the filaments aligns impressively with the direction of stretching when we look at the orientation projection maps (Fig. 2c). This result is even more striking when we quantify the three-dimensional orientation of the filaments over a 120 *µ*m thick portion of the aggregate (allowing more than 10 cell layers to be imaged). We define both an azimuthal angle *θ* with respect to the stretching direction and a polar angle *φ* with respect to the viewing direction (Fig. 2d). If the filaments were arranged to form a helix, they would appear as a ring in this stereoplot, whereas if arranged parallel, a dot would appear. On day 1, small peaks are visible in stretched aggregates at *θ* equals to 90^*°*^, corresponding to tilted filaments at the surface (Fig. 2b,e) while the actin filament orientation is almost uniform in the CTL spheroids. At day 2, we observe a remarkable peaked distribution for *θ* centred around 0 in the stretched aggregates (Fig. 2e) while no preferential orientation is observed in the CTL spheroids. In stretched aggregates, elongated cells orient along the direction of stretch in the inner part of the aggregate, initially having a perpendicularly tilted orientation on the surface (Fig. 2a-b) with an angle that decreases with time (Fig. 2c-e). The magnetic stretcher thus drives the 3D alignment of the actin filaments in the direction of stretching in the core of the aggregate.

### Stretching drives myogenesis

Actin orientation is not the only clue to muscle cell differentiation (38). The expression of myogenic factors or the one of proteins required for muscle contractile function, are generally used to quantify differentiation level. These proteins and factors have a temporal pattern of expression (38). Early, intermediate and mature muscle specific differentiation markers can be distinguished (39). Myogenesis is schematically regulated by myogenic factors from myogenic differentiation factor 1 (Myod1) and Myogenin (Myog) to myogenic factor 6 (Myf6) (40). Moreover, myosin heavy chain isoforms (Myh), troponin (Tnnt) and creatinin kinase M-type (Ckm) are specific markers of mature muscle. The expression of specific myogenic genes: Myod1 (early differentiation factor), Myog (intermediate differentiation factor) and Myf6 (late differentiation factor) or Myh, Tnnt and Ckm (specific markers of mature muscle) was measured by quantitative PCR and compared with the CTL spheroid on day 1. The measurement showed an increase in all myogenic genes after 3 days of differentiation (Fig. 3a). This shows that a purely cellular 3D environment is suitable for myogenesis. The increase in mRNA expression even counteracts the slight decrease due to magnetic labelling observed in 2D at short time scales (Fig. S2b). Furthermore, if there is an improvement in myogenesis in 3D, the most striking result is that this phenomenon is even enhanced by mechanical stimuli. Indeed, the mRNA expression levels of each myogenic gene are increased by stretch stimulation, with expression levels at least 1.4 times higher in stretched aggregates than in spheroids on day 3 especially for specific markers of mature muscle.

To ensure myogenic differentiation at the protein level, the expression of proteins required for muscle contractile function must also be determined. Myogenic factor 6 (Myf6) and fast myosin heavy chain (Myh4) proteins were measured by Western blot analysis. After 3 days of differentiation in the stretched condition, expression of both Myh4 and Myf6 proteins was increased compared to the non-stretched spheroid at day 3 (Fig. 3b). The expression of the late myogenic protein troponin was measured by immunofluorescence. The expression of this protein was much higher in stretched aggregates after 3 days of differentiation than in non-stretched spheroids at the same time (Fig. 3c).

**Fig. 3.**
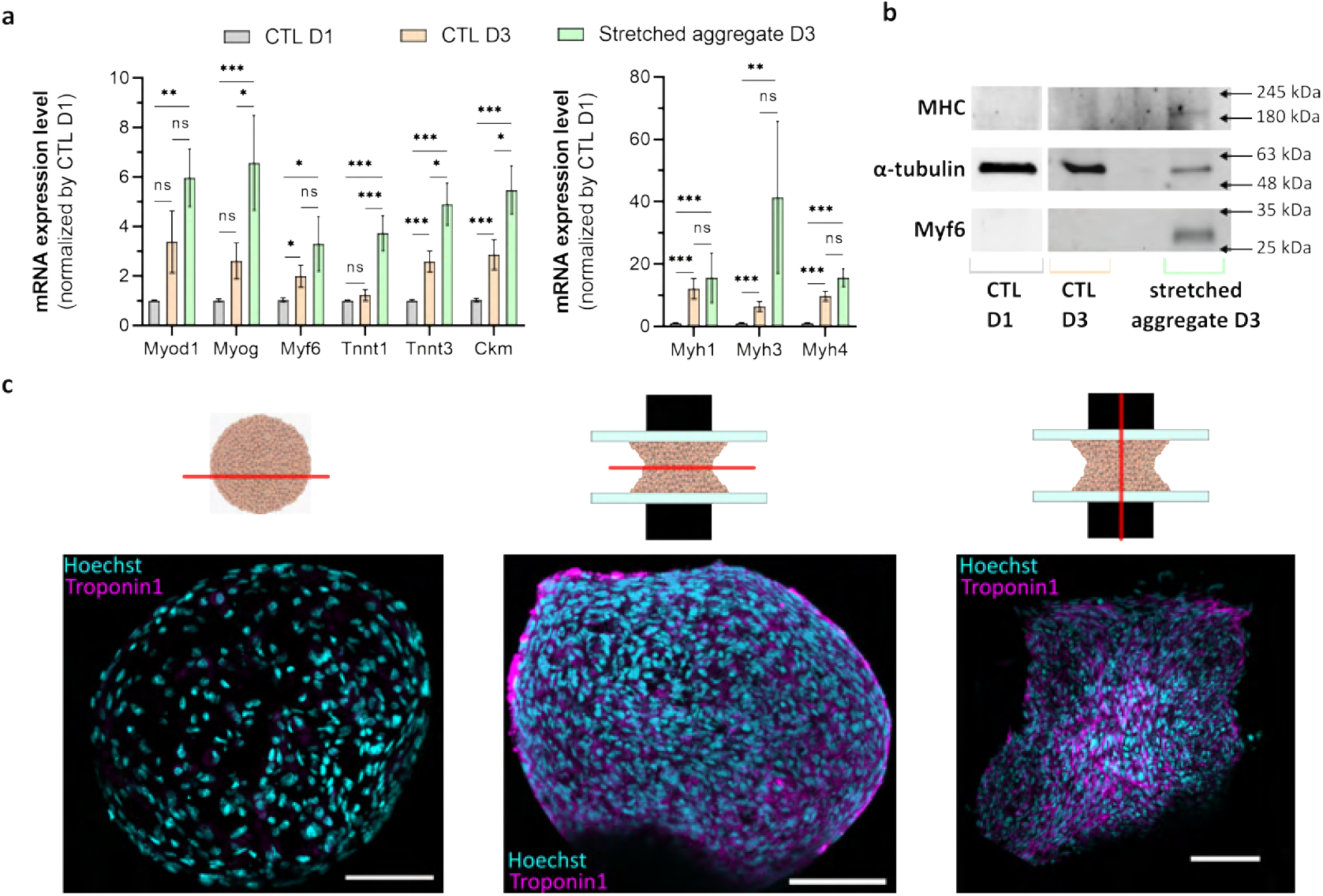
Myogenic differentiation enhanced by stretching. **a)** Myogenic marker expression levels, determined by quantitative reverse transcription polymerase chain reaction (RT-qPCR). Relative mRNA expression levels are measured for control spheroids at D1 and D3, and for stretched aggregates at D3. RPLP0 was used as a housekeeping gene. At least N = 6 for control spheroids with 4 independent experiments, N=4 to 8 in 4 to 8 independent experiments for stretched aggregates at D3 for each gene. Mean *±* SEM. Myod1, myogenic differentiation 1, Myog, myogenin, Myf6, myogenic factor 6, Tnnt, troponin T, Ckm, creatin kinase M-type, Myh, myosin heavy chain; mRNA, messenger RNA; D, day. **b)** Western Blot of control spheroids at D1 and D3 and stretched aggregates at D3 for fast myosin heavy chain, Myh4 and myogenic factor 6, Myf6. *α*-tubulin is used as a loading control. **c)** Representative confocal images of a spheroid and stretched aggregates at D3. Immunostaining of troponin 1 protein (magenta) is superimposed with nuclei (cyan) in a transverse cryosection of a control spheroid (left), of a stretched aggregate (middle) and in a longitudinal cryosection of a stretched aggregate. Scalebar = 200 µm.

Thus, physical stimuli (both dimensionality and mechanical constraint) enhance overall myogenesis, but how this couples with the multicellular architecture and its long-range orientational order remains to be elucidated.

### Spatially resolved muscle differentiation patterns under stretching constraints

In the context of spindle-shaped cells, differentiation is generally associated with integer topological defects (20, 23). In stretched aggregates, the only topological defects clearly identified are at the surfaces: +1/2 defects are present on the lateral surface of the aggregates, but they tend to annihilate, whereas defects accumulate in the regions in contact with the slides (Fig. 4a). The expression of skeletal muscle troponin (Tnnt1), a specific marker of mature muscle differentiation, was studied by imaging mRNA using single-molecule fluorescent *in situ* hybridisation (41–43) on day 2 to look more closely at the pattern emergence (Myh1 is expressed in the same cells as Tnnt1 and gives the same differentiation pattern - Fig. S6). mRNA expression is maximal on the lateral surface of the aggregates but no specific effect can be associated with +1/2 defects and at the central part of the aggregate (Fig. 4b), correlating with the area of high defects density and possible integer defects. This result is also evident in the immunofluorescence images of the protein (Fig. 3c).

**Fig. 4.**
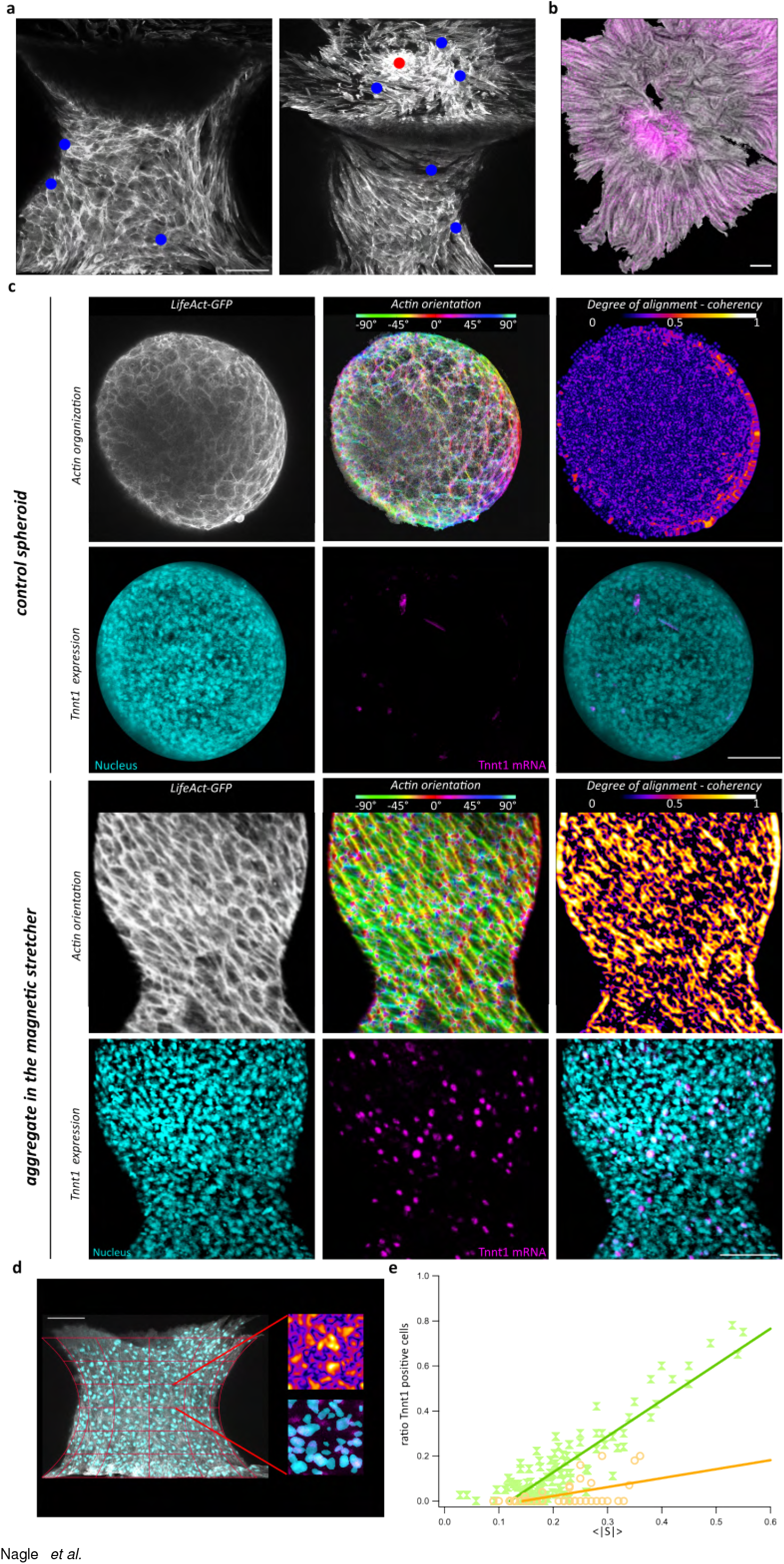
Differentiation pattern in stretched aggregates. **a)** 3D Reconstruction from confocal images of a stretched aggregates. Actin filaments are made visible using SiR-actin labelling. Topological defects are highlighted in blue (half integer defects) and in red (integer defect). Scale bar = 100 µm. **b)** Representative confocal image of troponin expression (magenta) surimposed with contour integration of actin filaments labelled with phalloidin. The section is on the attached part of the aggregate. Scale bar = 100 µm. **c)** Representative confocal images of sections of a spheroid and a stretched aggregate after 2 days. For each, on the first row, projection of actin filaments at the surface (left), local orientation of the filaments (middle) and the extracted coherency (right) are shown. On the second row, nuclei labelled with DAPI (cyan) and mRNA FISH signals for Tnnt1 (magenta) are visible and superimposed. Scale bars are 100 *µ*m long.**d)** Schematics of the definition of the different areas on the surface to count both positive cells and to measure local coherency. In this image, F-actin is shown in gray and nuclei in cyan. Scale bar = 100 µm. **e)**. Ratio of Tnnt 1 positive cells as a function of the local amplitude of the nematic vector determined on the surface of both stretched aggregates (green) and spheroids (orange). Linear fits obtained for each data sets are superimposed with the corresponding colors. For stretched aggregates, *χ*^2^ = 0.6, for spheroids *χ*^2^ = 0.1. In both cases N=4 different aggregates were studied.

Together with the orientation map, mRNA expression shows that interfaces are key regions for differentiation. We therefore focus on these regions. By performing conformal projections of curved interfaces for both actin structures and mRNA expression, the tilted structure of actin filaments on the surface of stretched aggregates on day 2 is imaged. The tilt angle is distributed from 0 to 40^*°*^ depending on the experiment and the position over the aggregate. Multicellular spheroids show shorter actin filaments and are more disordered (Fig. 4c) than stretched aggregates. Unsurprisingly, Tnnt1 mRNA expression is much higher than in control spheroids. To quantify cell alignment, we introduce coherency map (Fig. 4c). Coherency is the local amplitude of the nematic order parameter defined as 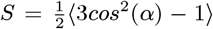 where *α* is the angle between filaments and the director, it equals 1 if the filaments are perfectly aligned and equals 0 in disordered orientation. By dividing the projected lateral surface into 50 *µ*m long squares (Fig. 4d), we examined the fraction of cells expressing Tnnt1 in correlation with the mean amplitude of the nematic order parameter *S* (Fig. 4e). The more aligned the actin filaments, the greater the number of myoblasts entering muscle differentiation, both for stretched aggregates and spheroids. However, the correlation is stronger when mechanical stress is applied. In the case of the stretched aggregates, a higher slope of the linear fit is measured with a larger *χ*^2^ value around 0.6. This indicates a higher mechanosensitivity under constraint.

## Discussion

The use of magnetically labelled cells allows the formation of cohesive multicellular structures that can be manipulated using micromagnets. These 3D tissues can be stretched at will as the micromagnets act as clamps at both ends, distributing the stretch uniformly throughout the volume (27) (Fig. 1). Applied to spindle-shaped cells such as mouse muscle precursor cells C2C12, this model mechanical stretcher gives rise to complex self-organizing structures.

Actin organization supports the cell and is a key element of its cytoskeleton. In addition, as myoblasts differentiate, supracellular actin fibres align with the body axis of the aligned muscle cells. So given the nematic properties of actin and its constitutive relationship with muscle structure (20), multicellular organization is seen from actin observation. First, cell layering occurs in these 3D systems when they formed. Surface effects have previously been observed in multicellular spheroids (37) or embryos (44) and have been associated with differential adhesion. In spheroids, differential cell tension and adhesion tends to flatten cells at the periphery with long actin filaments, while cells in the centre are rounder presenting cortical actin (37). In contrast, in stretched multicellular aggregates, on day 1, cells are all elongated. The overall structure of the actin filaments is not modified from the surface to the core but their global orientation twists from the surface, where it is tilted relative to the stretching axis, to the core, where it is not.

A multilayered organization of spindle-shaped cells has already been observed in organs and is tissue specific (45): whereas corneal cells show no correlation between successive layers, muscles have perpendicular crisscross layering (20, 46). This crisscross layerings can also be observed *in vitro* in bilayers systems (47). In muscle, these organizations persist for complex functions that may actuate organ formation (48). In our model system driven by external forces, the structure obtained one day after 3D myoblast tissue formation recapitulates this perpendicular organization (Figure 2b) at day 1. However, it is gradually modified by the application of external forces and subsequent rearrangements to provide a more globally oriented structure (Figure 2e) by decreasing the tilt angle at the surface. The still persistent tilt at the end of maturation could also be reminiscent of the one observed for cells covering polymer capillary bridges (17) except that in our case no chirality is visible. Surface tension may thus also be a factor explaining this multilayering. If not fully explained this configuration is like in the *in vivo* systems, it forms prior to muscle fibers orientation (48).

In spheroids, cells experience the same differentiation medium and could be driven towards the myogenic pathway. However, while actin filament length increases, global random organization persists. External mechanical forces are the actuators of cell alignment, they tend to elongate the actin filaments and the cells and orient them in the direction of stretch. C2C12 cells are well described as nematic cells (49), meaning that they tend to minimise possible bend and splay in 3D and thus align with each others (22, 50, 51). However, interface regions with glass slides tend to align cells parallel to the slides, potentially creating a tilt that is eventually gradually resolved depending on the nematic interaction and the actual Frank-Oseen elastic parameters, which may change during differentiation. Indeed, while confluent myoblasts show numerous defects and changes in orientation, myotube formation reduces this type of events and shows long-range ordering in 2D (Fig. S5).

Mechanical stretch not only drives the actin network cell alignment but it also enhances differentiation to mature muscle. Looking at both early (MyoD1), intermediate (Myog) and late (Myf6) factors of myogenesis (52) as well as mature muscle specific markers (Myh, Tnnt, Ckm), if 3D conditions are sufficient to induce muscle differentiation (Fig. 3a), differentiation is greatly improved by mechanical stretching, especially for mature muscle specific markers as the troponin isoforms. Contrary to scaffold approaches (53), purely cellular 3D culture conditions actually enhance muscle cell differentiation compared to 2D culture conditions. Upregulation levels obtained two days after differentiation medium addition and linear stretching are comparable to those obtained after 7 days of differentiation on 2D slides (Fig. S2). In addition, in the case of stretched aggregates, a microtissue is obtained, which means that its geometry is closer to *in vivo*. Muscle differentiation has a multilevel interplay (54), cell elongation and orientation (29) seem to play key roles in fusion and differentiation processes (52). Mechanical stretching by inducing both orientation and elongation accelerate this process as observed on 2D substrates (55, 56).

Protein and mRNA imaging point at two main hotspots for differentiation. First, high defect density region situated at the central part of the stretched aggregates exhibits a high differentiation level as was already observed in confined C2C12 patterns (23). In numerous studies starting from cell monolayers, the emergence of a third dimension is specifically linked to the dynamics of topological defects at different scales and in different systems (22, 57). In deformable substrates, depending on the nematic elastic constants and the boundary conditions, the emergence of disclinations is accompanied by deformation of the surface (58). For example, shape-changing vesicles encapsulating a microtubule-kinesin film form fibrillar protrusions to relieve the stress arising from half-integer defects (59). This recapitulates the mechanisms involved in the growth and regeneration of *Hydra* tentacles (60). In addition, *Hydra* head regeneration depends on integer topological defects (46), as does the formation of multicellular protrusions in confined myoblasts (23) or germ band extension in *Drosophila* (61). In stretched aggregates, defects relieve the stress induced by boundary effects due to cells parallel to the glass slide at the interface and create a cell structure parallel to the stretching direction in the center.

In addition, the lateral surfaces show higher differentiation. Specific surface tension forces are known to play a role in embryonic morphogenesis (62). Similarly, in stretched aggregates, surface cells experience maximum tension, which may explain their privileged differentiation. In addition, in myoblast-derived tissues, differentiation level is associated with a higher local orientational order (63) (Fig. S5). But if this tendency is observed both in spheroids and in stretched aggregates, the sensitivity associated with it, that is, the degree of correlation and the subsequent differentiation level obtained at a given order range, is higher when the aggregate is mechanically stimulated (Fig. 4e).

This study illustrates the complex interplay between physical cues and the emergence of differentiation maps in a 3D purely cellular complex system. It recapitulates the main features of myogenic processes and highlights the impact of mechanical constraints on the resulting structure but also on the sensitivity of the process. Indeed, mechanical constraints drive both cell alignment and actin orientation, but also a higher subsequent differentiation in long-range oriented regions. The ability of this tool to drive the formation of complex structures highlights the importance of linking cell signaling with cell organization. Mechanically-induced spatial reorganization modifies the local signals that each cell receives in a tissue, leading to the emergence of differentiation patterns. Mechanical stimuli enable the rational assembly of multicellular architectures and their potential remodeling. Combined with the modularity of synthetic cells (64, 65), this approach should potentially offer valuable insights into the dynamics of complex systems and facilitate the engineering of multi-layered structures. Overall, the mechanical stimulation of purely cellular 3D tissues provides powerful capabilities for cell and tissue engineering, as well as for the systematic study of multicellular organization. It also sheds light on the dynamics of complex systems and nematic tissues, demonstrating its potential to engineer customized self-organizing tissues or materials. The emerging understanding of morphogenesis through topology not only solves a key puzzle in biology, but also provides a mechanistic framework for future approaches to tissue engineering by highlighting processes with high robustness.

## Materials and Methods

### Cell culture and cell line construction (Lifeact-GFP)

C2C12 cells were obtained from ATCC (CRL-1772). The C2C12 Lifeact-GFP stable cell line was constructed by transfecting the cells with a Lifeact-GFP Puromycin plasmid (pLVX-LifeAct-GFPtag2 gift from Sylvie Coscoy) using Lipofectamine3000 reagents (L3000001, Invitrogen). Cells around 50% confluency in 6-well plates were incubated with a 1:1 DNA to Lipofectamine 3000 Reagent ratio for 24 hours. Cells were then selected with puromycin and sorted by FACS (Fluorescence-Activated Cell Sorting FACS Aria Fusion, BD Biosciences). The 10% of cells with the highest fluorescence were dispensed at one cell per well in 96-well plates to obtain individual clone cell lines after amplification. After 2-3 weeks of amplification, all the clones were expressing a similar level of Lifeact-GFP but only a few were differentiating. The clone selected presented a similar differentiation phenotype to C2C12 wild-type cells. C2C12 Lifeact-GFP cells were cultured in complete medium corresponding to Dulbecco’s modified Eagle’s medium (DMEM, Gibco), supplemented with 1% Penicillin-Streptomycin (P/S, Gibco), 2 *µ*g/mL puromycin (P7255, Sigma) and 10% Fetal Bovine Serum (FBS, Gibco).

### Citrated iron oxide nanoparticles

Maghemite superparamagnetic citrated iron oxide nanoparticles (*γ* − *Fe*_2_*O*_3_) were obtained by alkaline coprecipitation of Fe(II) chloride and Fe(III) chloride according to Massart’s procedure (66) followed by an oxidation thanks to the addition of boiling iron nitrate and finally the addition of sodium citrate. The adsorption of citrate anions to the nanoparticles ensures the electrostatic stability of the suspension. The resulting solution is composed of nanoparticles of about 8 nm diameter.

### Cell labelling

C2C12 cells were magnetically labelled by an incubation with a solution of iron oxide nanoparticles at [Fe] = 1 - 2 mM and 5 mM citrate in RPMI 1640 medium (Gibco) for 30 minutes leading to an uptake of about 3 pg of iron per cell (measured by magnetophoresis). Iron oxide nanoparticles (NP) are incorporated through the endocytosis pathway and are stored in the endosomes as described in Rivière et al. (32).

No influence of the iron oxide NP uptake was observed on the cell viability, the metabolic activity, or the ability to differentiate (27, 67, 68).

The magnetization of the multicellular aggregates stays almost constant during 3 days, nanoparticles are not eliminated from cells (Fig. S3c).

### Metabolic activity

Cell metabolic activity was quantified thanks to a fluorescent resazurin-based assay (TOX8-1KT, Sigma-Aldrich). The assay was performed 2 hours (day 0) or 24 hours after nanoparticle uptake (day 1). The resazurin-based solution was incubated 2 hours at 1:10 in DMEM (without phenol red) according to the supplier’s protocol. Fluorescence was measured for each condition thanks to a plate reader (Enspire, Perkin Elmer) with an excitation wavelength at 560 nm and an emission wavelength at 590 nm in a 96-well plate.

### Immunofluorescence analysis on 2D samples

Cells in 2D were fixed for 15-20 min in 4% paraformaldehyde (PFA) at room temperature (RT). They were then permeabilized 15-20 min in 0.1% Triton X-100 at RT and non-specific interactions were prevented by an incubation with 5% BSA (#05479, Sigma-Aldrich) for 1h at RT. Myh4 was labelled thanks to the MF20 mouse primary antibody (dilution 1:200 in 0.5% BSA D-PBS 1x, DSHB) incubated for 2h at RT. Cells were then incubated for 2h at RT with a goat anti-mouse Atto550 antibody (dilution 1:500 in 0.5% BSA D-PBS 1x, 43394, Sigma). Nuclei were stained with Hoechst 33342 (dilution 1:1000 in D-PBS 1x, H3570, Invitrogen) for 15-20 min at RT. Images were obtained by confocal microscopy thanks to the LSM 780 Zeiss microscope equipped with a 20x water immersion objective (W Plan-Apochromat 20x/1.0 DIC, Zeiss). Represented images correspond to Z-projections on about 10 µm.

### RT-qPCR measurements

Total RNA was extracted from multicellular aggregates with the NucleoSpin RNA Kit (740955, Macherey-Nagel) following the manufacturer’s protocol. For aggregate disruption, each aggregate was rinsed with D-PBS 1x and resuspended in 350 µL lysis buffer and disrupted with about 30 one-second pulses performed with a Biovortexer and a spiral pestle (918034 and 918044, Biospec). Complementary DNAs were obtained with the SuperScript II Reverse Transcriptase kit (18064022, Thermofischer Scientific) and random hexamers (C1181, Promega) according to the manufacturer’s instructions. Quantitative PCR was performed with SYBR Green Master Mix (4309155, Thermofischer Scientific). The RPLP0 gene coding for the 60S acidic ribosomal protein P0 was used as a reference transcript. The primer sequences used for RPLP0, Myod1, Myog, Myf6, Tnnt1, Tnnt3, Myh1, Myh3, Myh4 and Ckm are presented in Fig. S2a.

### Protein extraction and Western Blot

For protein extraction, multicellular aggregates were rinsed with D-PBS 1x and put in an ice cold solution of 30 mM Tris-EDTA (pH adjusted with concentrated HCl to 7.4), 1 mM DTT (dithiothreitol), 1x anti-phosphatase cocktail (Roche, 04906837001) and 1x anti-protease cocktail (Roche, 11836170001). Multicellular aggregates were disrupted using lysing beads (Precellys lysing kit, P000912-LYSK0, Bertin) and the Precellys24 tissue homogenizer device (Bertin). Samples were then left for 30 min in ice and centrifuged for 20 min at 12 000 g and 4°C. The supernatant was retrieved and protein concentration was quantified by Bradford assay.

The multicellular aggregate protein extracts were then used for Western Blot analysis. Proteins were separated on SDS-polyacrylamide 4-20% gels and transferred on PVDF membranes. After rinsing the membranes with TBS-Tween 20 buffer (TBST), the membranes were blocked for 1h in the EveryBlot blocking buffer (12010020, Biorad) and incubated overnight at 4°C with primary antibodies rabbit anti-Myf6 (sc-301, Santa Cruz Biotechnology), or mouse anti-fast myosin heavy chain (Myh4) (MF 20, DSHB) diluted at 1:1000 with 5% BSA. Membranes were then washed 3 times with TBST and incubated for 1h with the anti-mouse (#7076, Cell Signaling) or anti-rabbit (#7074, Cell Signaling) horseradish peroxidase-linked secondary antibodies diluted at 1:2000 in the Everyblot blocking buffer. Peroxidase activity was revealed using a chemiluminescent detection kit (Amersham ECL Prime Western Blotting Detection Reagent, RPN2232, Cytiva) and the Syngene PXi device (Ozyme). Alpha-tubulin was used as the loading control and immuno-probed with the mouse anti-alpha tubulin antibody (T5168, Sigma).

### A. Biocompatibility of nanoparticles

The biocompatibility of citrated iron oxide NP incorporation for mouse muscle precursor cells C2C12 was carefully assessed by measuring the metabolic activity and the differentiation level.

Measurements on the metabolic activity of cells show no difference in comparison with control cells 2 hours after NP incorporation at day 0 and at day 1 (Fig. S1a). Besides, differentiation tests were performed after NP incorporation showing no phenotypic difference on the myotubes formation. Fig. S1b shows representative images of cells after 7 days of differentiation. Both in the control condition and with the incorporation of nanoparticles at day 0, long myotubes are obtained. These elongated polynucleated cells express myosin heavy chain, a marker of myogenic differentiation involved in the contraction of functional muscular cells, in both control and NP conditions after 3 days of differentiation (Fig. S1c).

Finally, NP incorporation decreases slightly mRNA levels for all genes related to myogenic differentiation at day 1 after NP incorporation, probably due to the upregulation of genes involved in NP uptake and storage such as ferritin (69). However, the same mRNA levels of the myogenic marker genes are recovered with or without NP incorporation for most genes at day 3 and for all genes at day 7 of differentiation (Fig. S2). In conclusion, NP uptake does not alter significantly the cell metabolic activity or the ability of cell to differentiate at medium (3 days) and long term (7 days).

### Formation of C2C12 spheroids

C2C12 control spheroids were obtained as described in (37). In summary, C2C12 cells grown in 2D and at 70 − 80% confluency were incubated for 30 min with a solution of iron oxide nanoparticles at [Fe]=2 mM supplemented with 5 mM citrate. After 2h of incubation in complete medium, cells were trypsinized, centrifuged and resuspended in a minimal volume (a few hundreds of µL). Suspended cells where then attracted in spherical agarose molds of 1.2 mm diameter, previously coated for 30 min with an anti-adhesive rinsing solution (07010, Stemcell Technologies), with permanent magnets placed below each mold. After an overnight incubation at 37°C, 5% *CO*_2_, spheroids were removed from the molds gentle pipetting of the surrounding medium.

### The magnetic stretcher device

The magnetic stretcher device is composed of a tank with two identical 3D printed polymer holders (PA2200, polyamide) in which a permanent neodymium magnets (6 × 6 mm, Supermagnete) is inserted and put into contact with a soft iron tip of 400 *µ*m diameter. Cells are deposited on each microtip protected by a 100 µm thick glass slide. One of the magnetic pieces is fixed while the second one can be moved thanks to a step-by-step motor (Z812B, Thorlabs) actuated by a controller (TDC001 or KDC101, Thorlabs). Temperature is regulated at 37°C. A homemade cover was built to enable the displacement of the objective while keeping the box closed and allowing the flow of 5% *CO*_2_, 95% humidity in the chamber (Okolab controller). Simulations of the magnetic field and the magnetic field gradient between the two magnets are performed using the software COMSOL Multiphysics (additional module AC/DC). The volume magnetic force *f* applied to the cells is proportional to the magnetic field gradient, since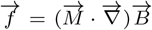, where 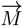 is the volume magnetic moment and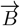 is the magnetic field.

### Aggregate formation in the magnetic stretcher

After tank sterilization, the glass slides on each micromagnet are incubated with Matrigel (356234, Corning) diluted at 0.5 mg/mL in cold DMEM for 1h at room temperature for coating. After rinsing, the tank is filled with DMEM supplemented with 10% FBS, 1% P/S, 2.5 ug/mL Amphotericin B, 2 µg/mL puromycin and 1x anti-oxidant supplement (A1345, Sigma).

C2C12 cells are incubated for 30 min with a solution of iron oxide nanoparticles at [Fe]=1-2 mM supplemented with 5 mM citrate. After 2h of incubation in complete medium, cells are resuspended at 5 million cells/mL in complete medium. 10 µL of suspended cell solution (equivalent to 50k cells) are deposited on each micromagnet forming a hemisphere of non cohesive cells. The two micromagnets are then approached at a distance of 350 to 400 µm as shown Figure 1d-e. After an overnight incubation, the tank medium is changed for the differentiation medium composed of DMEM with 2% HS, 1% P/S, 2.5 ug/mL Amphotericin B and 1x anti-oxidant supplement. A linear stretch at 1 µm every 3 minutes of the cohesive multicellular aggregate is started for 6 hours and the mobile piece is reapproached of about 40 µm at the end of the stretching. The medium is changed with fresh differentiation medium every 24 hours for the next two days.

### Live bright field and 2-photon microscopy imaging

Bright field images were obtained with a Leica DMIRB inverted microscope and a 10x objective. Live fluorescence images through 2-photon microscopy were obtained with an SP8 Leica microscope coupled with an Insight DeepSee tunable laser. Acquisition were obtained at a wavelength of 980 nm with a 20x water immersion objective (HCX APO L 20x/1.00W, Leica)

### Imaging of fixed 3D samples

Control spheroids or stretched aggregates were fixed for 1h at RT in 4% PFA. Stretched aggregates were fixed *in situ* then retrieved with a micropipette. 3D samples were permeabilized and non-specific interactions were blocked by incubating the 3D samples in 5% BSA and 0.5% Triton X-100 for at least 4h at RT. SPY555-actin or SiR-actine (Spirochrome) was incubated with the samples for at least 24 hours at a 1:1000 dilution in D-PBS 1x to ensure an optimal actin imaging. Samples were then imaged by confocal microscopy thanks to the LSM 780 Zeiss microscope and with a 20x water immersion objective (W Plan-Apochromat 20x/1.0 DIC, Zeiss).

### Actin filament orientation extraction

To obtain actin filament position and length, ridge detection filter was applied. Actin filament orientation was extracted thanks to Orientationpy the derived 3D pythonic version of the well-known ImageJ plug-in OrientationJ (Biomedical Imaging Group, Ecole Polytechnique Federale de Lausanne, Switzerland) (70) from either live 2-photon microscopy images or confocal images on fixed samples. Orientation of actin filaments was assessed from images located from 0 to 120 µm below the surface of the spheroid or the stretched aggregate. The representative images were obtained from ImageJ plugin OrientationJ with a gaussian gradient.

### Single molecule fluorescence *in situ* hybridization

mRNA FISH probes were purchased from ACDBio (Mm-Myh1-O2 REF:539381, Mm-TnnT1-C3 REF:466911). Fixation and labelling protocols were optimized from published protocols (43, 71). Briefly, multicellular aggregates were fixed *in situ* for 1h, in a solution of 4% PFA, 0.25% glutaraldehyde, 0.1% Tween, 5mM EGTA, 0.2% Triton X-100, 1:100 Alexa Fluor™488 phalloidin (Molecular Probes). After fixation, they were incubated in PBS with 1:100 phalloidin 488, 0.1% Triton X-100, 1% BSA for 4 hours to ensure optimal retention of cytoskeleton structures.

Probes were heated to 40 °C for 10 min. Samples were then incubated in Eppendorf tubes overnight (O/N) in probes diluted 1:50 in Probe Diluant (REF300041, ACDBio). The next morning, samples were washed 3x 15 min in wash buffer under agitation at RT (RNAscope Wash Buffer Reagents, ACDBio, REF:310091). Samples were incubated with each amplification solutions consecutively for 35 min at 40*°*C and then washed 3 × 5 min at RT in wash buffer. Samples were incubated in HRP-C1 for 20 min at 40*°*C and then washed 3 × 5 min at RT in wash buffer. Samples were incubated in TSA-570 (REF:323272, ACDBio) or TSA-650 (REF:323272, ACDBio) diluted 1:1500 in TSA buffer for 35 min at 40*°*C and then washed 3x 5 min at RT in wash buffer. Samples were incubated in HRP blocker for 20 min at 40*°*C and then washed 3 × 5 min at RT in wash buffer. Samples were stained in DAPI (solution from the RNAscope kit) for 5 minutes or O/N in 1:1000 Spy505 (Spirochrome) and then mounted in ProLong™ Gold (Invitrogen, P36930) in between two coverslips (thickness 1.5mm, 22mm × 22mm) with double spacers (Thermo Scientific™ Gene Frame, 65 µL) between them.

### Image analysis

To conformally project aggregates surfaces from 3D volumes, we used both the ImageJ plugin LocalZ projector and the Matlab Deproj application (72) without noticing important changes as our samples are relatively flat over the 150 *µ*m height of the observation. Nuclei detection was performed using StarDist plugin. Coherency is extracted from ImageJ plugin OrientationJ in 2D images and from Python orientationpy plugin in 3D reconstruction. Line integral contour of cells is obtained using MatLab developped program.

## ACKNOWLEDGMENTS

This work was supported by the Program Emergence(s) de la Ville de Paris (Grant MAGIC Project) and the Île-de-France Region via the DIM BioConvS. The study was supported by the Labex Who Am I?, Labex ANR-11-LABX-0071, the Université de Paris, Idex ANR-18-IDEX-0001 funded by the French Government through its Investments for the Future program and the French Defense Procurement Agency (DGA-AID) France. This project has received financial support from the CNRS through the Tremplin Action. We acknowledge the ImagoSeine core facility of the Institute Jacques Monod (member of the France BioImaging, ANR-10-INBS-04) and especially Nicolas Valentin on the FACS platform. We thank the staff of the MPBT (physical properties – low temperature) platform of Sorbonne Université for their support. We acknowledge Veronique Thevenet (Laboratoire MSC, UMR 7059, Paris) and Aude Michel (PHENIX, UMR 8234, Paris) for providing us with the nanoparticles.

---

**Fig. S1.**
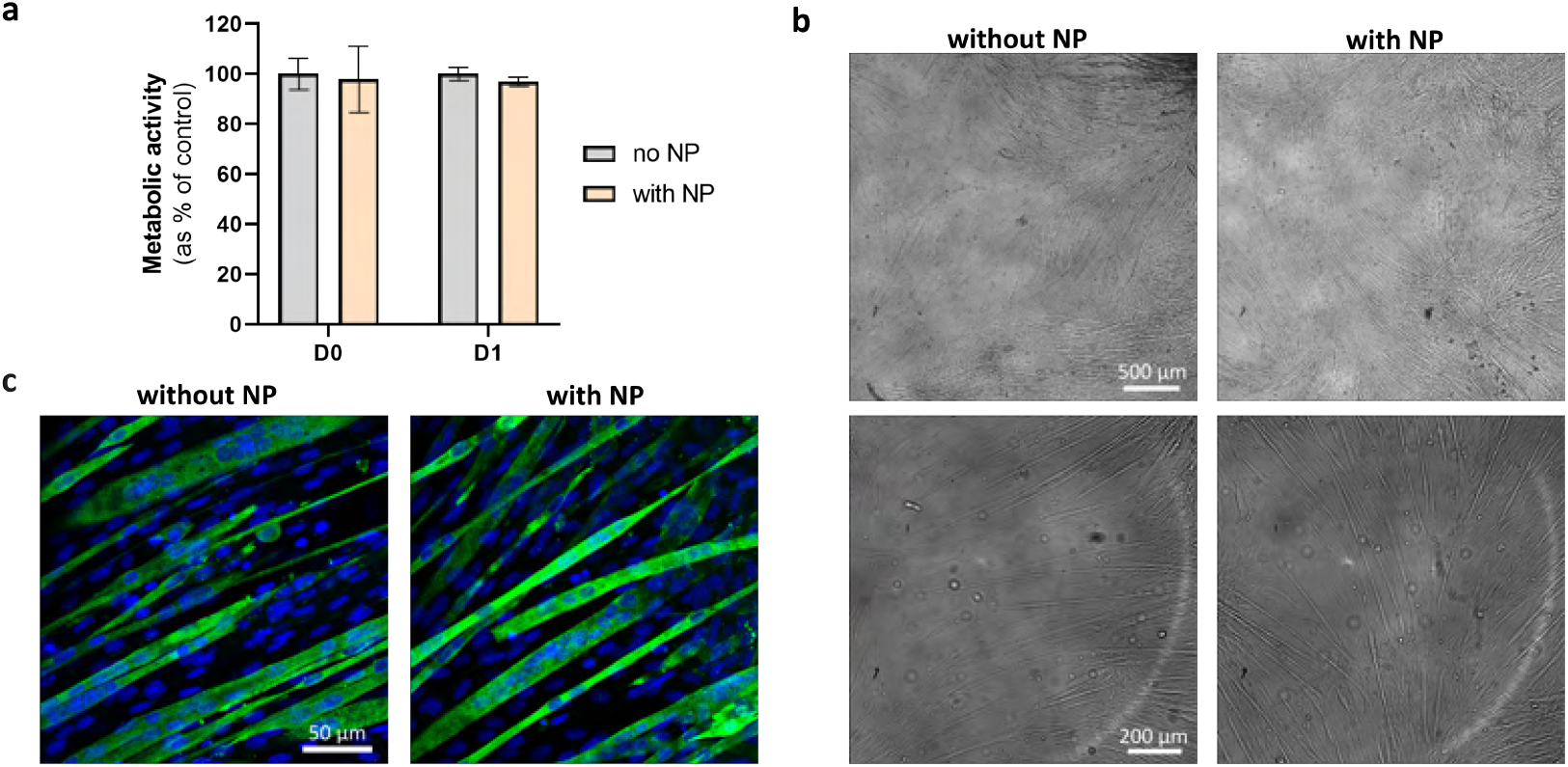
Nanoparticle incorporation in myoblasts does not impair their metabolic activity or their ability to differentiate. **a)** Metabolic activity of C2C12 cells with or without nanoparticle (NP) 2 hours (D0) or one day (D1) after the incorporation ([Fe] = 2 mM for 30 min). Mean +/-SEM. **b)** Representative images of C2C12 after 7 days of differentiation with or without NP incorporation at day 0. Long myotubes are visible in the entire field of view in both conditions. **c)** Immunofluorescence representative images of C2C12 after 3 days of differentiation and incorporation of NP at day 0. Myh4 and nuclei are shown in green and blue respectively.

**Fig. S2.**
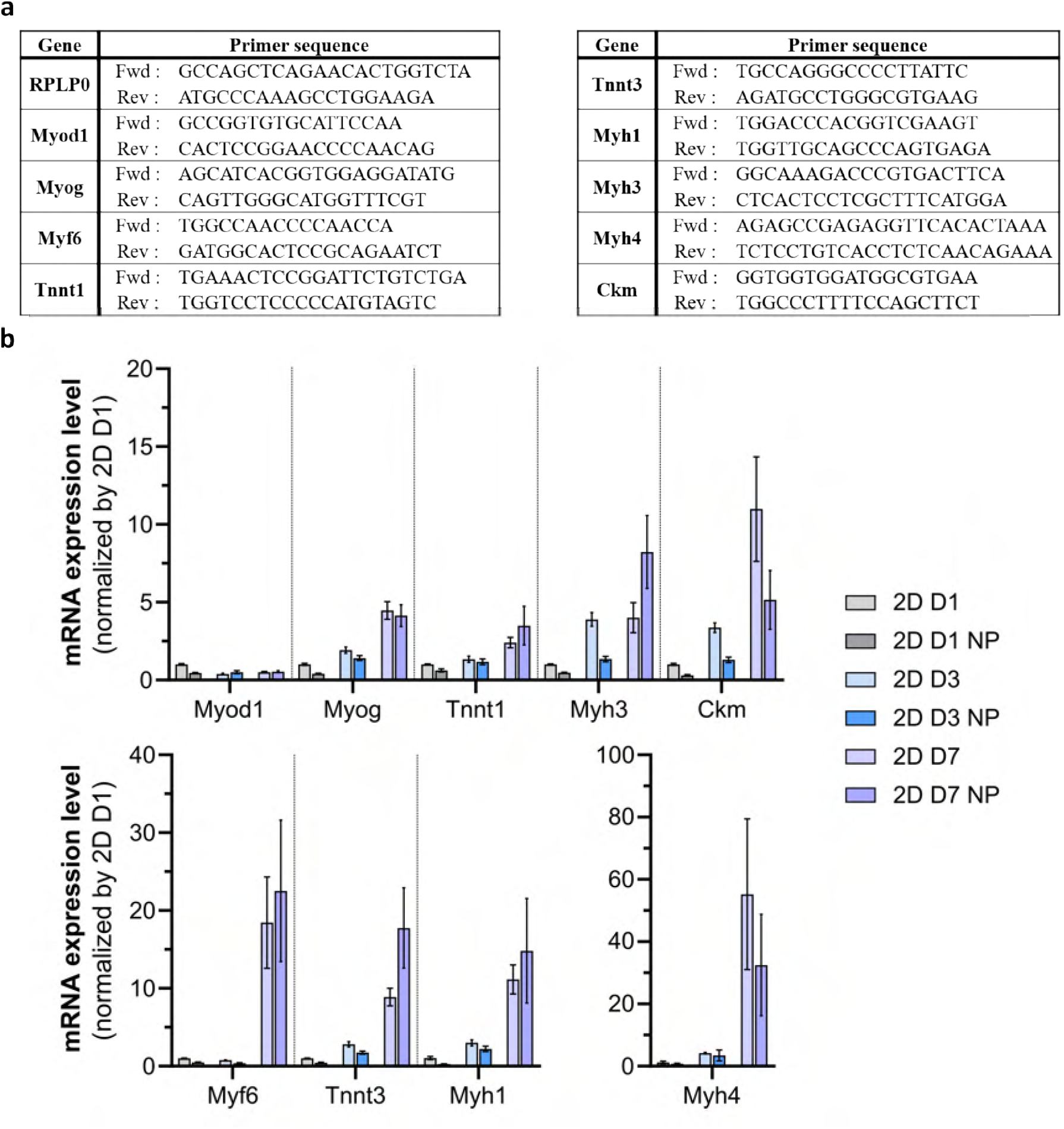
mRNA expression levels measured by RT-PCR for 2D cultures with or without NP incorporation and after 1, 3 or 7 days of differentiation. **a)** Primer sequences used for RPLP0, Myod1, Myog, Myf6, Tnnt1, Tnnt3, Myh1, Myh3, Myh4 and Ckm. **b)** Expression levels are normalised by the mRNA levels at day 1 without NP incorporation (2D D1). RPLP0 was used as a housekeeping gene. Mean +/-SEM (N=4).

**Fig. S3.**
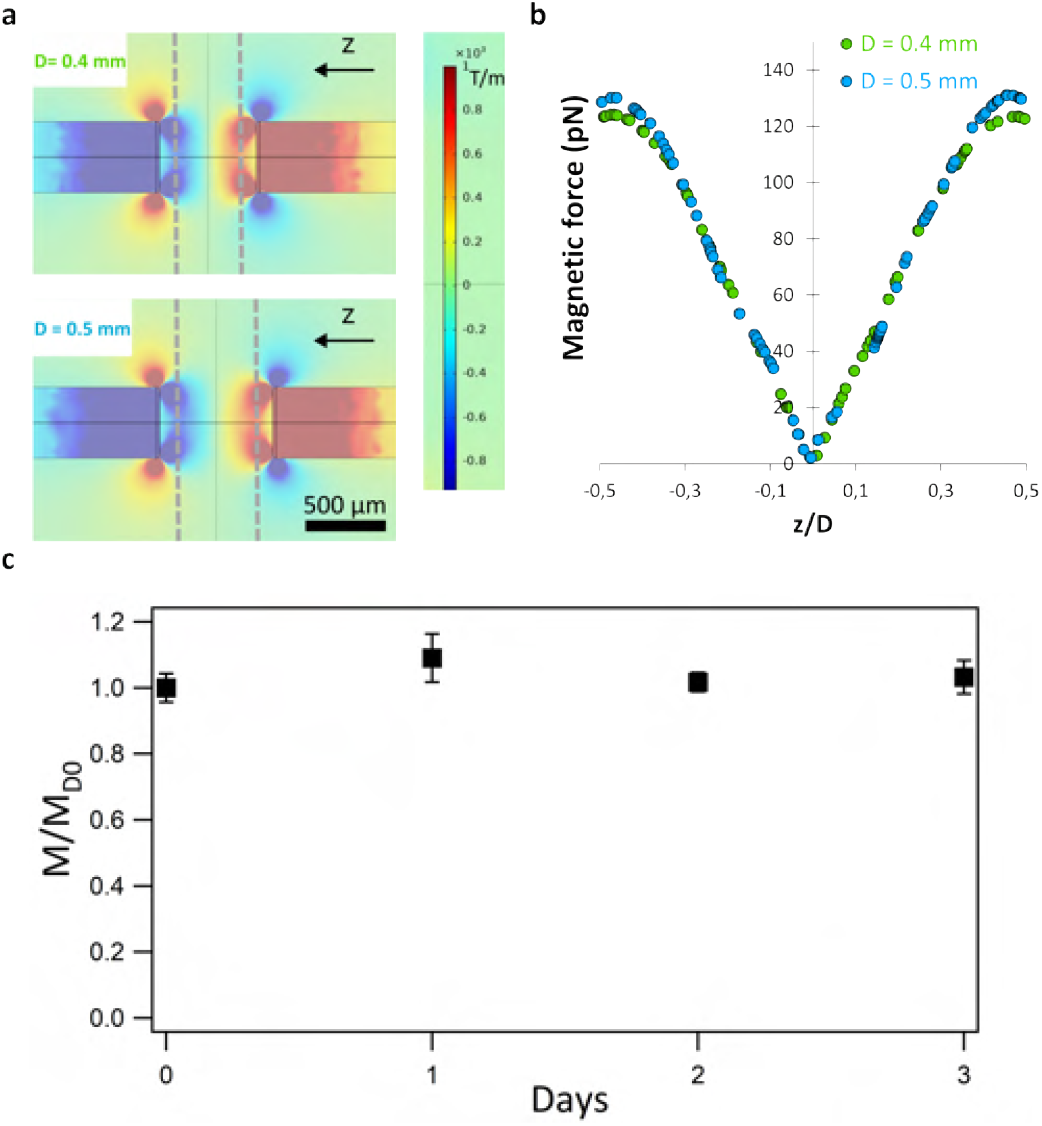
Magnetic forces exerted on magnetically labelled cells. **a)** Simulation of the magnetic field gradient in the magnetic stretcher for distances *D* of 0.4 mm (*D* before stretching) and 0.5 mm (*D* after stretching) between the two pieces.The positions of the glass slides are materialised by grey dashed lines. **b)** Simulated magnetic volume forces between the two micromagnets as a function of the position *z* normalised by interglass slide distance *D* for distances *D* of 0.4 mm and 0.5 mm. The two profiles are very close. **c)** Evolution of the magnetization of magnetic CTL spheroids over a week. The total magnetization of multicellular aggregates is compared between 0 to 3 days for aggregates of comparable initial size. Mean *±* SEM are indicated. N=5 independent experiments for each point.

**Fig. S4.**
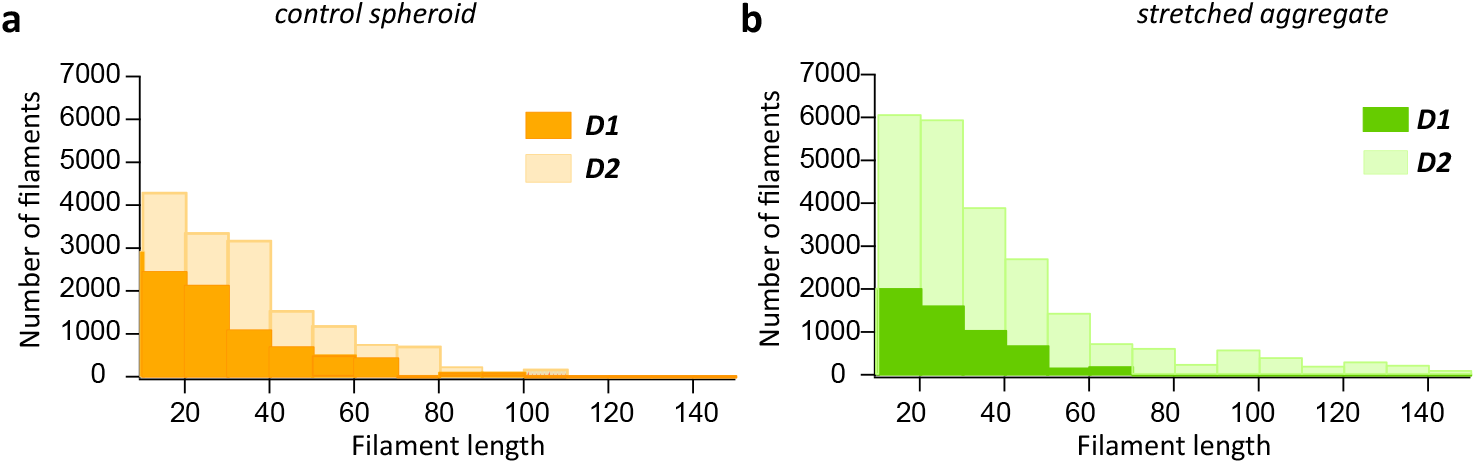
Evolution of the actin filament length in spheroids and in stretched aggregates. **a)** Distribution of filaments detected above 10 µm in length for control spheroids on D1 or D2 (N=4 independent experiments). The number of filaments increases but their distribution does not change. **b)** Distribution of filaments detected above 10 µm in length for stretched aggregates on D1 and D2 (N=4 independent experiments). The number of filaments increases as well as the number of long filaments.

**Fig. S5.**
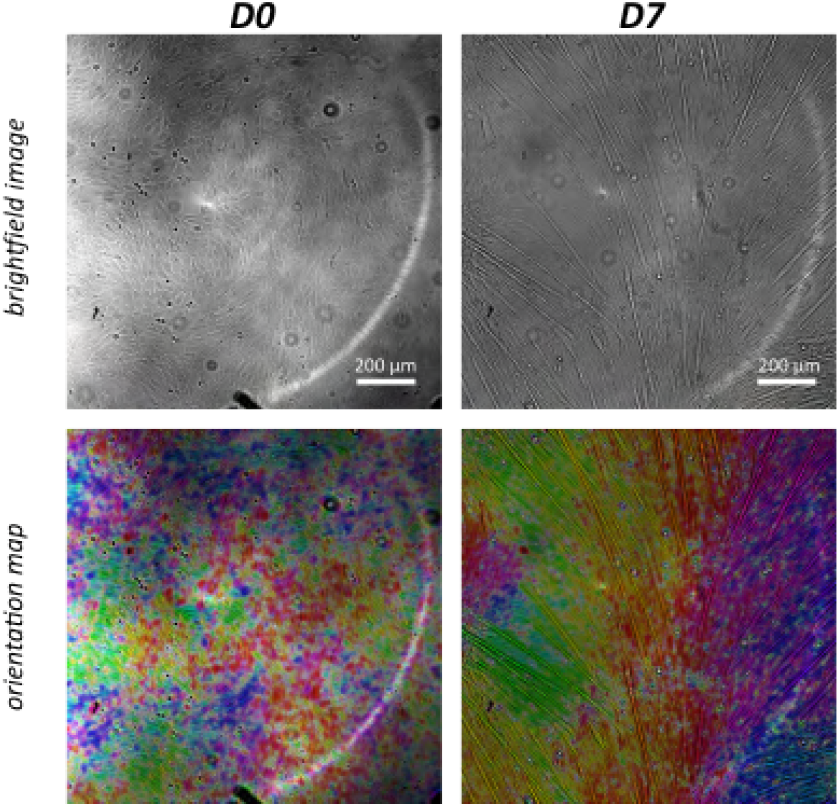
Evolution of the cell orientation during myogenesis. Myoblasts plated on 2D slides at confluency at day 1 (left) are compared with differentiated myoblasts after 7 days under differentiation medium (right). Representative brightfield images (top) and superposition with orientation map (bottom). On the orientation map, the brightness is proportional with the coherency. Scalebar = 100 µm.

**Fig. S6.**
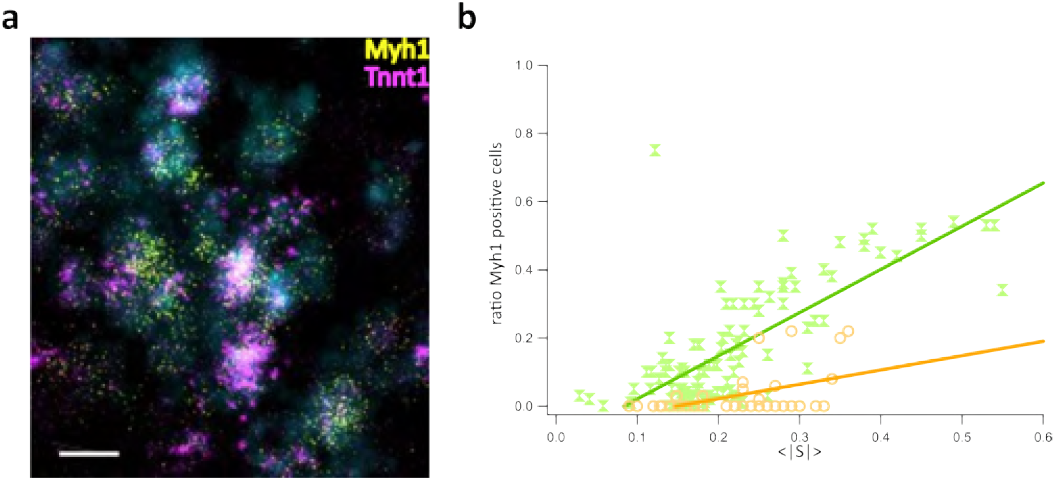
Evolution of the coherency during myogenesis. **a)**. Representative confocal image of cells on the surface of stretched aggregates labelled with mRNA FISH probes for Tnnt1 (purple) and Myh1 (yellow). Nuclei are superimposed in cyan. Scale bar = 10µm **b)**. Ratio of Myh 1 positive cells as a function of the local amplitude of the nematic vector *S* determined on the surface of both stretched aggregates (green) and spheroids (orange). Linear fits obtained for each data sets are superimposed with the corresponding colors. For stretched aggregates, *χ*^2^ = 0.4, for spheroids *χ*^2^ = 0.05. In both cases N=5 different aggregates were studied.

